# Compression of protein secondary structures enables ultra-fast and accurate structure searching

**DOI:** 10.1101/2025.09.12.675776

**Authors:** Runfeng Lin, Sebastian E. Ahnert

## Abstract

Protein structure prediction has undergone a revolution with the advent of AI-based algorithms, such as AlphaFold and RoseTTAFold. As a result, over 200 million predicted protein structures have been published. This wealth of structural data has created a need for rapid structure comparison algorithms, such as Foldseek, which enable efficient searches across this vast space of protein structures. Here we introduce a new ultra-compact representation of protein structure in the form of Secondary Structure Elements (SSEs). These are short sequences around 8% of the length and with 10% of the information content of full amino acid sequences and 3Di sequences. We show that, despite this compression factor, SSEs can be used as a highly effective tertiary structure comparison tool, with accuracy that approaches that of Foldseek, while offering a 200-fold speedup. In addition SSEs offer comparable performance to Foldseek in domain boundary retrieval. Furthermore we show that the particular way in which SSEs encode structure can also be used to specifically detect proteins that differ due to conformational change. These findings demonstrate that SSEs offer a valuable complementary approach for protein structure characterisation.

## 1 Introduction

Proteins are fundamental components of every living cell, and their functions are closely tied to their three-dimensional structures. In the course of biological evolution, protein sequences can diverge substantially but still result in highly similar folded structures. Multi-Sequence Alignments (MSA) reveal that proteins with as little as 20% sequence identity [1] can yield near-identical structures. Widely used databases such as Pfam use hidden Markov models to create MSAs of amino acid sequences. [2] in order to classify protein families and assign domains.

Recent advances in artificial intelligence (AI), such as AlphaFold [3] and RoseTTAfold [4] have greatly increased the number of predicted protein structures to around 200 million. However, with the rapid growth of tertiary structure databases, direct structure comparison tools like TM-align [5] or Dali [6] are too slow to scan the entire database. To address this, methods like Foldseek (FS) [7] have been developed. Foldseek uses machine learning to convert 3D protein structures into 3Di sequences and applies MMseq2 [8] to efficiently filter out unrelated structures. This approach is both fast and highly accurate.

Protein secondary structures can by itself already provide valuable insights into protein tertiary structure [9, 10, 11, 12]. Proteins with very different sequences but similar folds have been identified using detailed secondary structure alignments [10]. These methods have been further improved by refining the classification of secondary structures [12] and the development of a Monte Carlo model that combined secondary structure and hydrogen bonding information in order to narrow down the possible ways a protein can fold [11, 12].

Based on the close relationship between protein secondary and tertiary structures we introduce a compact sequence representation of protein structure in the form of Secondary Structure Elements (SSEs), and show that this encoding enables ultra-fast and accurate searches in the space of protein tertiary structure. The full pipeline is shown in Figure 1. We first extract the secondary structure information of a given protein and encode it as a sequence of characters, each of which encodes the type (helix or sheet) and length of a secondary structure element. These secondary structure element (SSE) sequences are compared using a highly customized Smith-Waterman algorithm [13], which incorporates a substitution matrix and element-specific gap penalties. These gap penalties and substitution matrices are optimized through genetic algorithms to recover SCOPe [14] domain boundaries, making the alignment approach more adaptable compared to standard sequence alignment methods that use uniform gap penalties. The tertiary structure similarity (in form of a TM-score) of aligned sub-SSEs sub-strings is predicted using a hybrid ResNet-BiLSTM network, trained on randomly selected pairwise comparisons of 70% of SCOPe proteins. The SSE sequences are around 8% of the length of the full amino acid sequence, and of the 3Di sequence used by Foldseek as a representation of tertiary structure (see Figure 2). This results in a significantly smaller sequence space (by a factor of ≈ 14.75^−*L*^) and an approximately 10-fold compression of the information content compared to conventional amino acid sequences and 3Di sequences. This in turn facilitates protein structure searches that are up to 200 times faster than Foldseek with accuracies that approach those of Foldseek. Moreover, as we show below, SSEs can also identify proteins that differ due to conformational changes. These results demonstrate the potential of SSE-based approaches for the rapid characterisation and analysis of protein tertiary structure.

**Figure 1.**
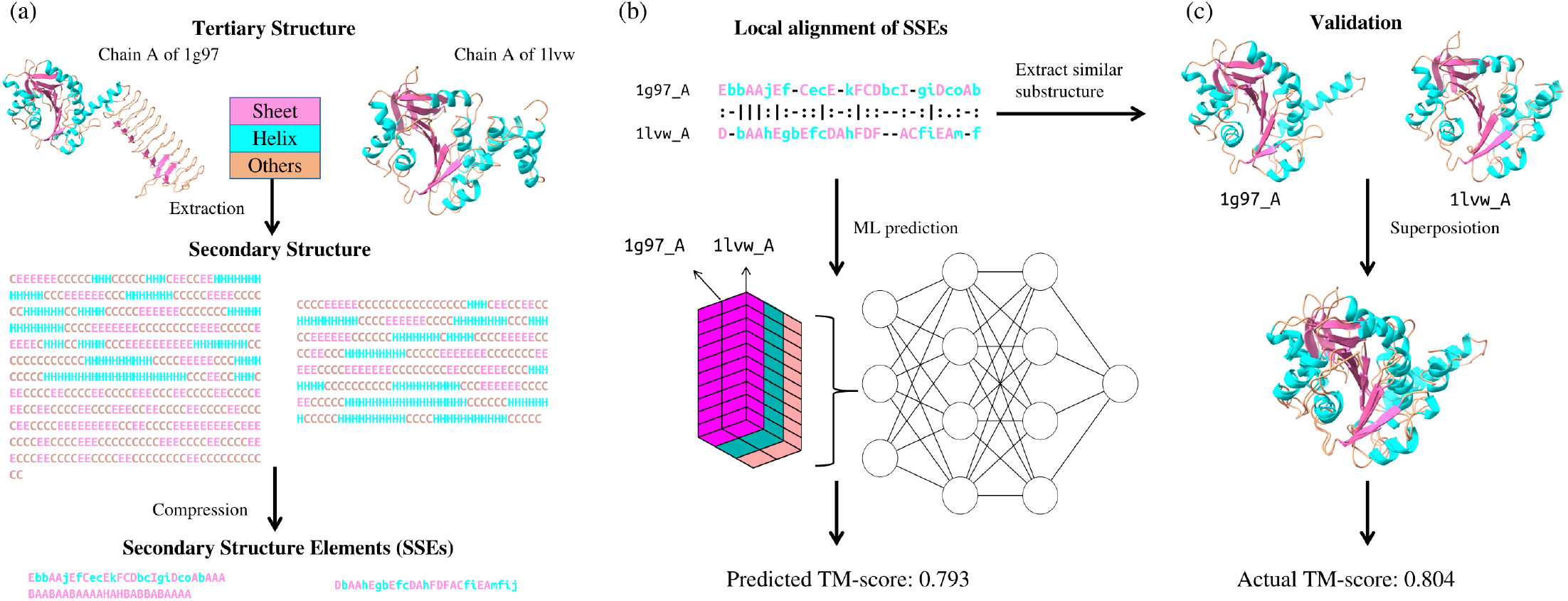
SSE-Based Structural Comparison Pipeline. (a) Structural processing pipeline illustrated using protein pair 1g97 A (*Streptococcus pneumoniae*) and 1lwv A (*Thermotoga maritima*), showing tertiary structure conversion to secondary structure elements (SSEs). (b) Local SSE alignment identifies conserved substructures (TM-score=0.804) despite low sequence identity (18.8%). (c) Machine learning prediction (TM-score=0.793) using alignment-derived features closely matches experimental validation, demonstrating the framework’s reliability.

**Figure 2.**
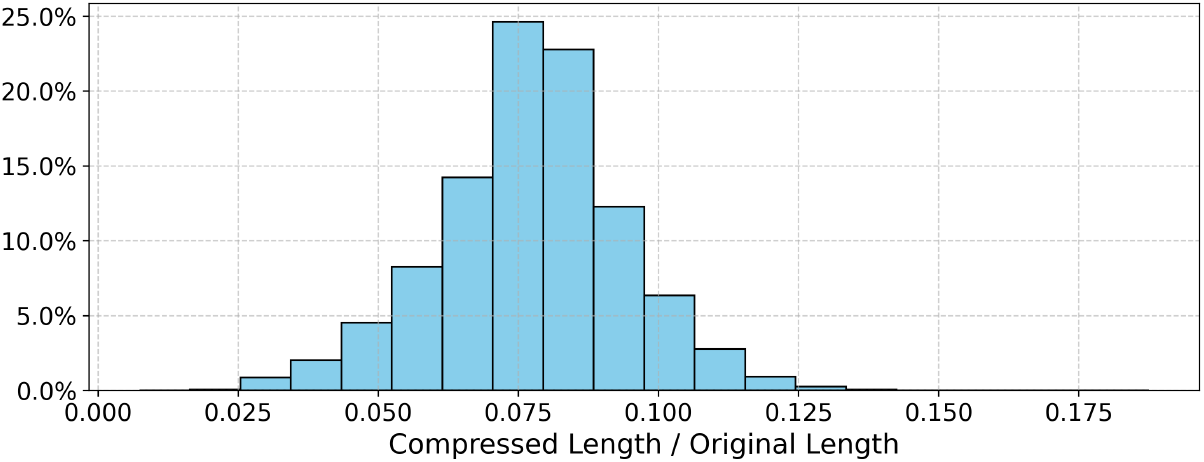
Compression Ratio of SSEs which has a mean ratio of 0.0774.

## 2 Results

### 2.1 SSEs offer substantial speedups over structure comparison methods based on full-length sequences

We benchmarked all-to-all alignments of 1000 proteins (i.e. 1e6 comparisons) on a single CPU core. The CPU-only SSEs pipeline (no filtration) completed in 770 s, of which 710 s were spent on similarity prediction. Introducing pre-filtration reduces the total runtime to 46 s (6 s for alignment + 36 s for prediction), at 89% sensitivity. Offloading prediction to an RTX 4060 GPU yields 91 s total (32 s prediction) or 11.6 s with filtration (1.6 s prediction). This is more than 200 times faster than Foldseek [7], which required 2400 s (95% of pairs filtered) and BioPython’s local aligner of primary structure, which took 1720 s on the same dataset. Table 1 summarizes the resulting speedups of SSEs (CPU and GPU, with and without filtration) over Foldseek, primary-structure methods, and TM-align; all variants can be further accelerated via multi-core CPUs or more powerful GPUs.

**Table 1:**
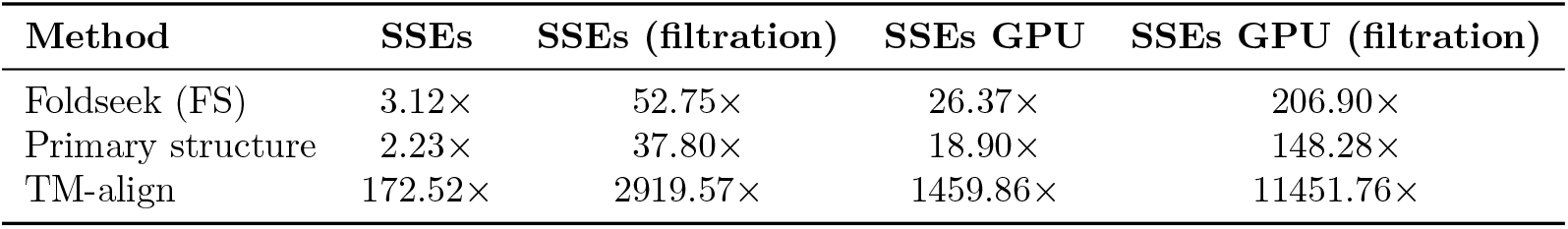
Speedup of SSE-based methods over other alignment approaches. Note that Foldseek employs pre-filtration of the data.

Although SSEs uses an alphabet of 52 characters—larger than the 20 standard amino acids and 3Di residues—the reduction in sequence length to around 8% significantly decreases the the-oretical search space by a factor of approximately 14.75^*L*^ (see Methods). According to Shannon entropy [15], the original sequence contains *L* · log_2_(20) bits of information, while the SSE representation encodes ≈ 0.08*L* · log_2_(52) bits, leading to a compression efficiency of approximately 9.5. This substantial reduction in possibility space not only improves computational speed, but also enhances the scalability of the method for large-scale protein structure analysis.

### 2.2 SSEs offer accuracy that approaches that of structure comparison methods based on full-length sequences

The performance of the SSE-based structure comparison method was evaluated by comparing it to Foldseek (FS) [7] using proteins drawn from the SCOPe [14] database, which offers a comprehensive structural classification of protein domains. A subset of 18500 SCOPe proteins was randomly selected as a training set, with the constraint that sequence identities remain below 0.9 within this set. The randomly selected testing set also maintained sequence identities below 0.9 among its members, resulting in a final testing set of 7,929 proteins. As shown in Figure S3 (a), when tasked with identifying structurally similar proteins (defined here as pairs of proteins with TM-scores greater than 0.6) the SSE-based method, attains a maximum F1 score of 0.738, which is more than 80% of the maximum F1 score of 0.884, achieved by FS for the same task with full-length 3Di sequences.

The SSEs method identifies 83,625 pairs of proteins with TM-scores above 0.6, while Foldseek (FS) extracts 132,859 pairs—approximately 58.9% more than SSEs. Notably, nearly all pairs detected by SSEs are also captured by FS, indicating that FS has high sensitivity within its detection range. However, FS identifies an additional 36,981 high-scoring pairs (TM-score *>* 0.6) that are either missed by SSEs or incorrectly extracted with TM-scores below 0.5. These FS-unique pairs are clearly distinct from structurally dissimilar cases. However, the 3306 similar pairs that are uniquely extracted by SSEs cannot be reliably distinguished from structurally dissimilar ones.

Further analysis reveals that most of the mismatches in SSEs arise from two main scenarios: (1) pairs with borderline TM-scores (0.6–0.65), where subtle structural similarities may not be well-represented by SSEs alone; and (2) pairs with very few SSEs, resulting in limited structural resolution. This limitation is partly due to the design of the SSE similarity matrix, which is trained to recover SCOPe domains typically consisting of at least five SSEs. When the number of SSEs falls below this threshold, the extracted features may not be sufficient to represent the full structural similarity (see Figure S1).

To provide further context, local sequence alignment of amino acids using Biopython aligner [16] was also benchmarked on this dataset, achieving a maximum F1 score of 0.306. Local sequence alignment identifies 60,526 pairs with TM-scores above 0.6, of which only 18,325 pairs (30.2% of its total extractions) fall within the TM-score range of 0.6–0.7. In comparison, the SSEs method extracts 37,542 pairs in this range (45% of its total), while FS identifies 76,817 pairs (57% of its total). These results demonstrate the significant advantage of the SSEs method over traditional sequence alignment in extracting and predicting similar substructures.

To ensure meaningful comparisons, all alignments were filtered to include only proteins with at least 50 residues, as more than 90% of proteins range from 50 to 200 amino acids [17]. Additionally, an extra filter requiring a minimum of five SSEs elements was applied to maintain robust comparisons.

Since both FS and SSEs were trained on the SCOPe database [14], we constructed an independent benchmark set to mitigate potential bias and fairly evaluate generalization performance. This set was derived from the RCSB Protein Data Bank (PDB) [18] using a stringent two-stage filtering protocol. First, to ensure internal diversity, we generated a non-redundant subset of the PDB by removing any protein with ≥90% sequence identity to another entry within the set. Second, to eliminate any overlap with the training data, we further filtered this subset by excluding any protein sharing ≥90% sequence identity with any entry in the SCOPe database. From this final collection of proteins, which was independent both internally and from our training data, 3,000 proteins were randomly selected to form the benchmark dataset.

As shown in Figure 3 (b), both FS and SSEs demonstrated improved performance on this unrelated dataset compared to their performance on SCOPe. The SSE-based method achieved a maximum F1 score of 0.821, which is 89% of the maximum F1 score of 0.915 achieved by FS. Using sequence identity alone achieved a significantly lower score of 0.69. In terms of pair extraction, FS identified 97,926 pairs with TM-scores above 0.6, while SSEs and local sequence alignment extracted 80,260 and 91,790 pairs, respectively.

**Figure 3.**
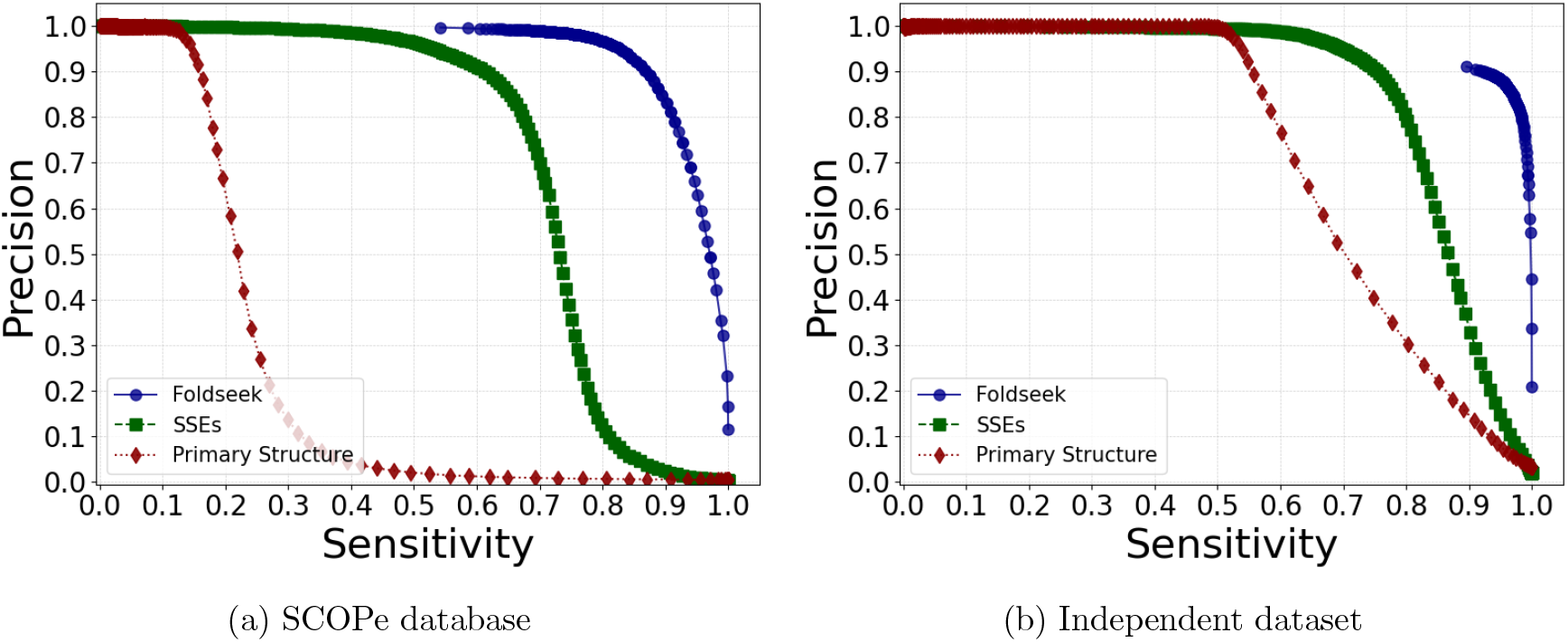
The accuracy of structure comparisons based on highly compressed SSE sequences approaches that of structure comparisons based on full-length sequences. Precision-sensitivity analysis on the (a) SCOPe benchmark dataset and (b) independent validation set with less than 90% sequence identity to SCOPe. FoldSeek (FS), using full-length 3Di sequences achieves maximum F1 scores of 0.884 (SCOPe) and 0.915 (independent). The SSE-based method, which uses a highly compressed representation (containing around 10× less information) achieves F1 scores around 80-90% as high (0.738 and 0.821). Local sequence alignment achieves much lower F1 scores (0.306 and 0.686).

### 2.3 SSEs enable the detection of conformational change in proteins

To understand why the SSE-based method extracts fewer pairs than the local sequence alignment method, we analyzed the distribution of extracted pairs (Figure 4). This analysis revealed that many pairs extracted by the SSEs method had high predicted TM-scores but only moderately high actual TM-scores. Upon closer inspection, we observed that these pairs often consisted of two or more similar substructures with distinct relative orientations. Typically, one half of each of these protein structures aligned well in 3D space, while the other half did not align at all, despite being structurally similar. This resulted in lower actual TM-scores. To address this, we designed a scanner to scan and split the corresponding protein pairs, identifying the best-aligned regions and normalizing the TM-score. This adjustment improved many of the lower TM-scores and increased the number of positive extractions to 89,513, and the corresponding F1-score to 0.837 (91% of the FS F1-score). Importantly this procedure allows us to specifically identify proteins that differ due to a conformational change, which could either represent two static states separated by evolution, or two different snapshots of very similar proteins that undergo dynamic conformational change in the cell.

**Figure 4.**
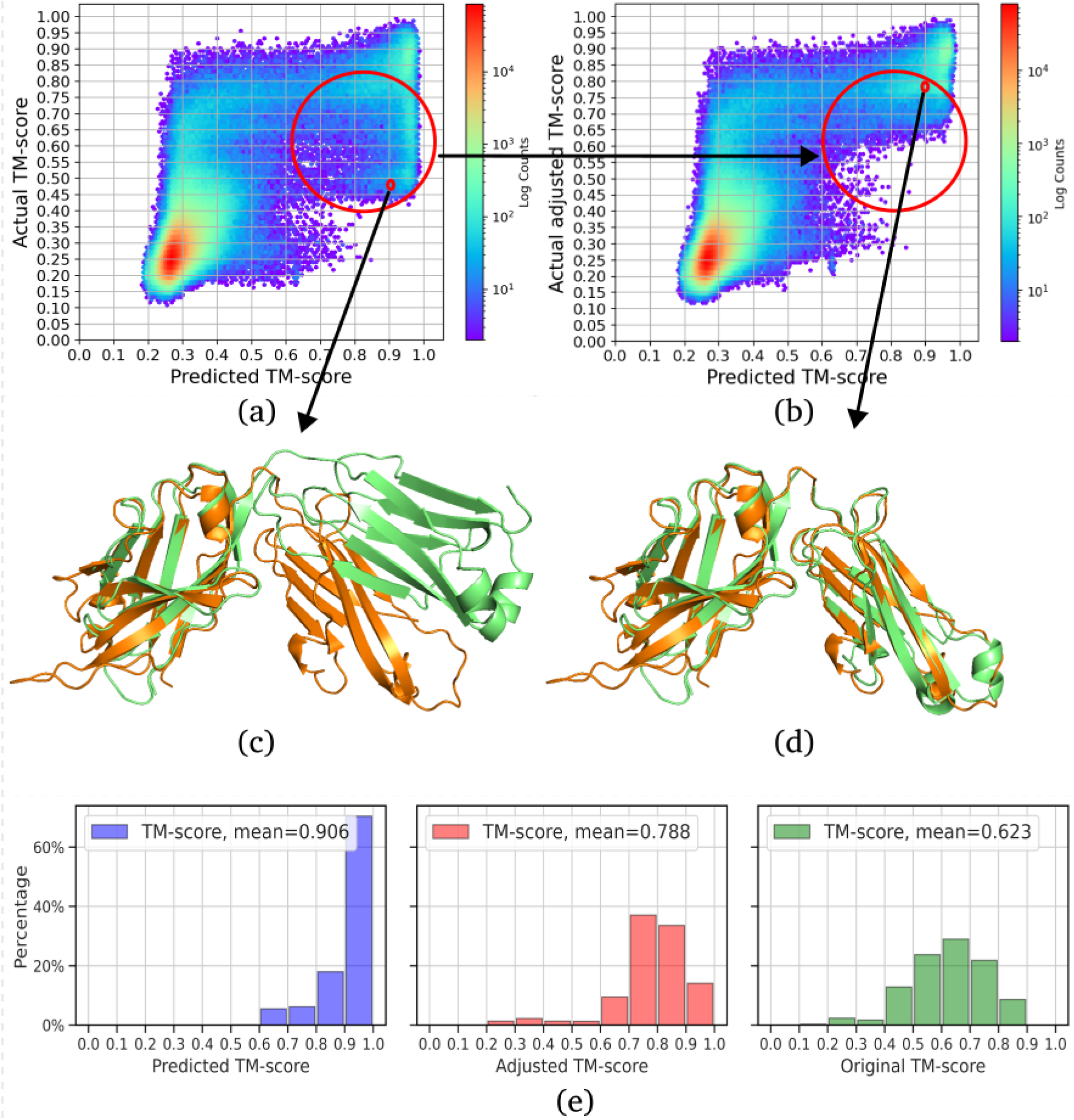
SSEs can detect conformational changes in proteins. (a) Correlation between SSE-predicted TM-scores and actual TM-scores. (b) Adjusted TM-score correlation after scanner-based normalization. (c) Structural superposition example from panel (b): 8dkf A (orange) and 1bfo C (green) exhibit 25.2% sequence identity in SSE-aligned regions and TM-score 0.499. (d) Enhanced alignment achieved by splitting 1bfo C (green) into two segments and re-superposing with 8dkf A (orange) with adjusted TM-score 0.794, demonstrating improved structural matching compared to panel (c). Their predicted TM-score is 0.912 based on SSEs. (e) Distribution of TM-scores for high predicted TM-score pairs with actual TM-scores, before and after adjustment. These figures show that SSE not only can predict similar structure but also detect conformational changes.

### 2.4 SSEs are highly effective at detecting similar tertiary structures with highly divergent primary sequences

TIM barrel proteins (like triose-phosphate isomerase) have very similar 3D structures even when their amino acid sequences are very different [19]. This structure is very common in enzymes — about 10% of enzymes use this shape. To test our method, we studied TIM barrel proteins from the CATH database (superfamily ID: 3.20.20.70) [20]. We filtered out sequences with over 90% identity to generate a dataset of 612 non-redundant structures. Initial analysis using local sequence alignment revealed a critical limitation: conventional sequence identity metrics failed to reliably distinguish structurally similar TIM barrel proteins (TM-score ≥ 0.6) from dissimilar ones, as demonstrated by the substantial overlap in their sequence identity distributions (Figure 5 (a)).

**Figure 5.**
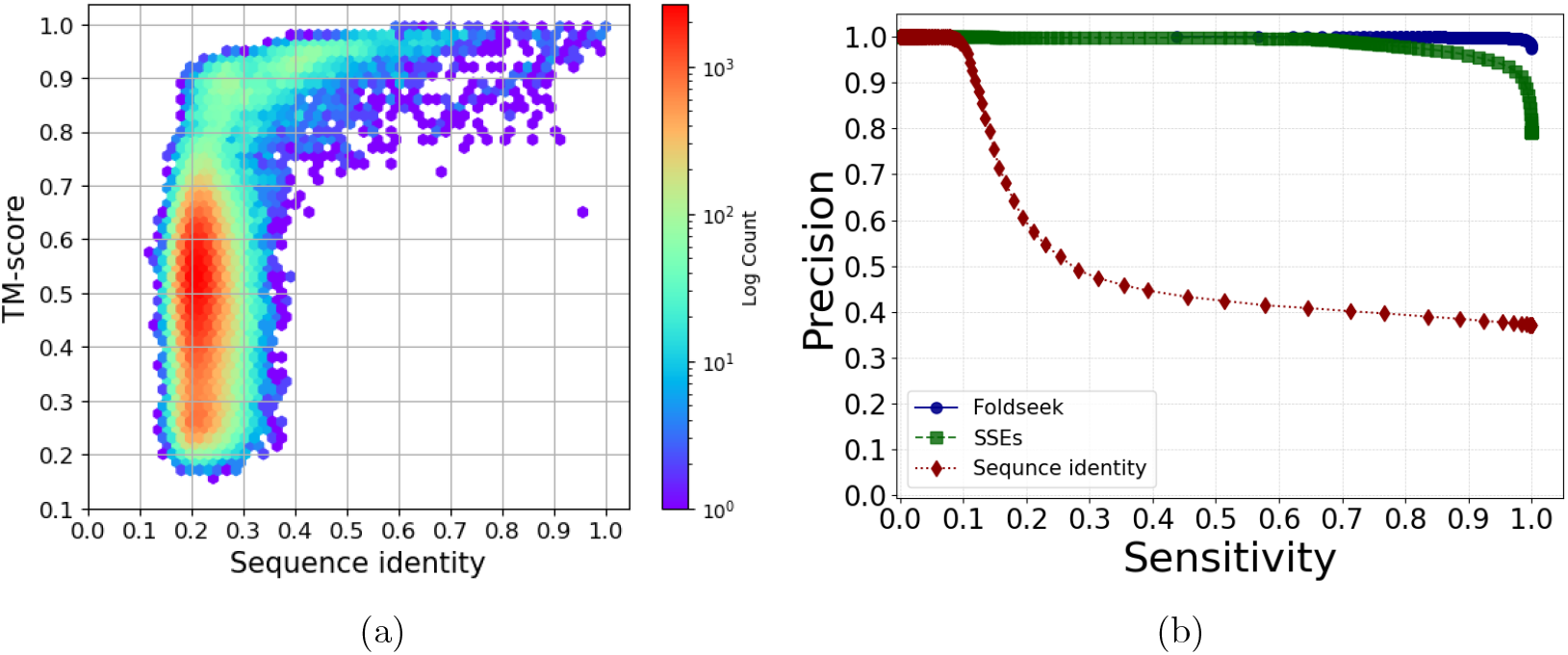
TIM Barrel Structural Comparison Analysis. (a) Correlation between local sequence identity and structural similarity (TM-score) in aligned regions. (b) Precision-recall curve analysis at TM-score threshold 0.6: FoldSeek (FS, F1=0.992) outperforms SSEs (F1=0.949) and traditional sequence alignment (F1=0.542) in detecting structural matches.

Both our SSE approach and FoldSeek (FS) showed better results than sequence alignment when proteins had low sequence similarity, as seen in Figure 5 (b). At TM-score ≥0.6 threshold, SSE detected 70,035 similar structure pairs (F1 score=0.949), which is much higher than the 33,057 pairs (F1=0.542) found by sequence alignment but lower than FS’s 118,647 pairs (F1=0.992). When we used a less strict cutoff (TM-score ≥0.5), SSE found 143,804 pairs compared to FS’s 175,497 pairs, while traditional sequence alignment only identified 90,020 pairs. Although FS performed best in both high-confidence matches (TM-score ≥0.6) and overall F1 scores, SSE still found many more matches than amino acids sequence alignment. These results confirm that SSE can effectively identify structural similarities that are hard to detect through sequence comparisons alone.

### 2.5 SSEs correctly identify domain boundaries, matching accuracy of methods based on full sequences

Although both Foldseek (FS) and the SSEs method confidently extract similar substructures, the biological significance of these substructures remains unclear. To address this, we investigated how well the boundaries of the extracted substructures align with domain boundaries from the SCOPe [14] and CATH [20] databases. For the CATH database, we excluded all proteins that also appear in the SCOPe database. Additionally, domains consisting of multiple disjoint regions were excluded to ensure clarity in boundary comparisons. For multi-domain proteins, we filtered out those with identical domains to avoid the extraction of full structures.

First, we evaluated domain recovery on a pairwise basis by comparing pairs identified by SCOPe and CATH as sharing domains. Since the SSEs method requires alignments to include at least five elements, we restricted our analysis to SCOPe and CATH domains containing five or more elements which excluded 1,879 from the initial 28,281 proteins.

The results show that the substructures extracted by both the SSEs method and Foldseek (FS) closely align with the boundaries defined by SCOPe and CATH. Although both methods were trained using SCOPe data, FS experienced a slightly more pronounced performance drop in the CATH database. In contrast, the SSEs method demonstrated slightly better performance in matching these boundaries compared to FS (see Figure 6).

**Figure 6.**
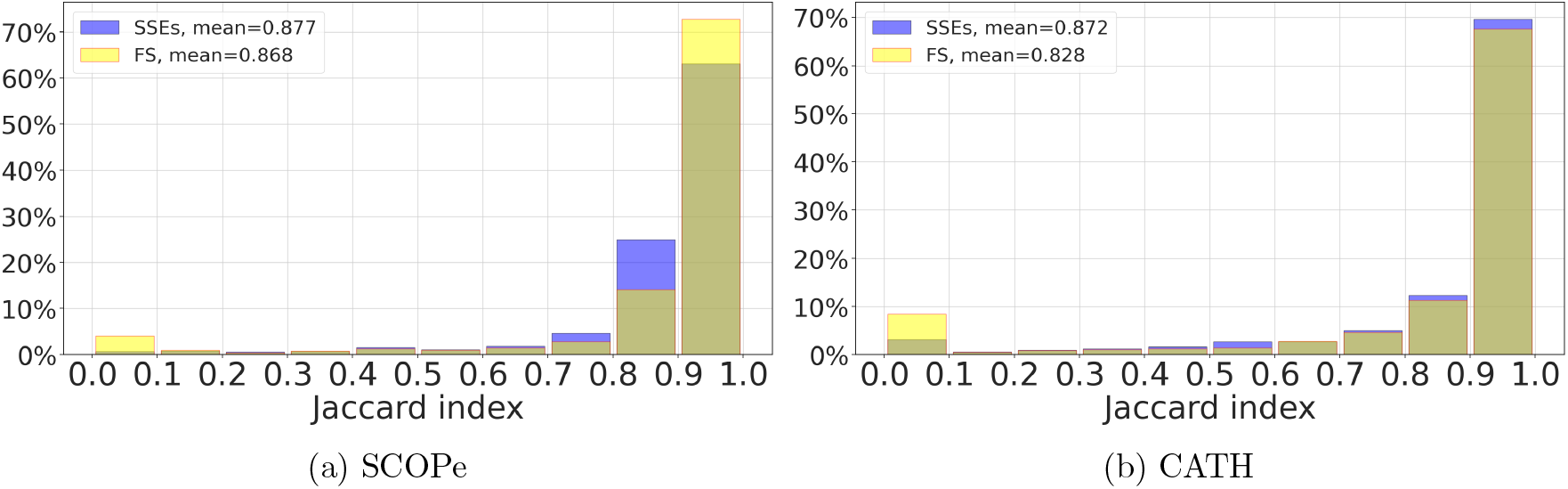
Domain Boundary Recovery Accuracy. (a) SCOPe and (b) CATH domain boundary alignment comparisons. Performance metrics show SSEs maintains consistent accuracy across databases (SCOPe: 0.872 vs CATH: 0.877) compared to FoldSeek (FS) (SCOPe: 0.868 vs CATH: 0.828).

However, the distribution of domain family sizes is highly skewed, which could introduce bias in pairwise comparisons of predefined pairs. To address this, we conducted a blind extraction and summarized the boundaries using Density-Based Spatial Clustering of Applications with Noise (DBSCAN). For this analysis, we selected substructures with a predicted TM-score above 0.5 for the SSEs method and a probability above 0.5 for Foldseek (FS). Overall, the SSEs method demonstrated superior performance in recovering full domains, while FS proved more effective at identifying boundaries that contain only part of the full structure (see Figure 7).

**Figure 7.**
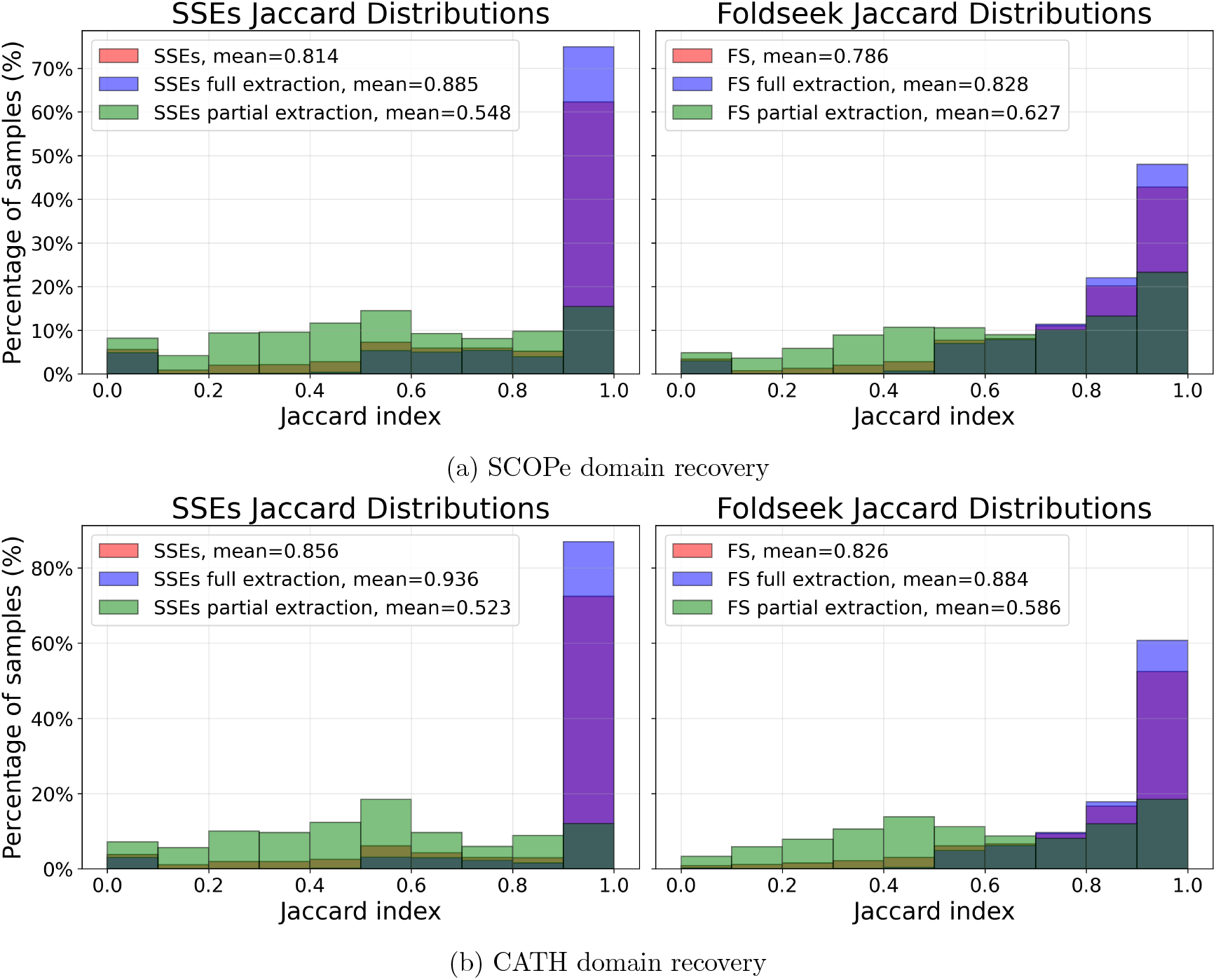
DBSCAN-Clustered Domain Boundary Recovery. (a) Jaccard similarity distribution between SSEs-predicted-reference SCOPe domains(left) and FoldSeek (FS)-predicted-reference SCOPe domains(right). (b) Jaccard similarity distribution between SSEs-predicted-reference CATH domains(left) and FoldSeek (FS)-predicted-reference CATH domains(right). SSEs demonstrate superior performance for complete domain recovery while FS shows better partial-domain detection, indicating complementary strengths in different biological contexts.

## 3 Discussion and Conclusion

The ten-fold compression of protein structure achieved using SSEs enables rapid structural comparisons. But the fact that tertiary structure comparisons are possible with SSEs is not just remarkable because of the drastically shortened sequence length. SSEs encode only secondary structure information. The fact that this is sufficient for accurate tertiary structure comparisons points to the way in which secondary structure severely constrains the configuration space of protein tertiary structure. The reliance of SSEs on secondary structure also explains why they are well-suited to detect conformational change, by detecting discrepancies between tertiary structure alignments and SSE comparisons.

We have shown here that SSEs provide a highly compact representation of protein structure that can be used for ultra-fast protein structure comparisons that are up to 200 times faster than established methods while also maintaining accuracy scores that are 80% or higher than those of established methods. Domain boundaries can be extracted with comparable accuracy. Moreover SSEs can be used to look for other structural hallmarks, such as conformational change. We also demonstrate, using the example of TIM-barrel proteins, that SSEs can identify similar protein structures even when their sequences are highly divergent.

These computational and biological capabilities suggest immediate applications in primary structural screening for drug discovery, evolutionary studies requiring rapid fold assessment, and tools for protein structure visualization.

Future development should prioritize three key areas: enhanced algorithms for multi-domain protein handling through dynamic element splitting, optimization of the compression-prediction pipeline to realize full theoretical speed advantages, and web server implementation with real-time visualization to enhance accessibility. Advancements in these directions could significantly expand our ability to decode protein structure-function relationships while maintaining computational efficiency, particularly for large-scale structural studies that encompass the entire universe of predicted protein structures.

## 4 Method

### 4.1 Secondary structure extraction and compression

Before studying the correspondence between SSEs and tertiary structure, the protein secondary structure needs to be assigned first. The assignment of secondary structures (SS) is carried out using the DSSP package under scheme 2. In this scheme, helices are represented by H and G; sheets are denoted by E and B, while the other elements are designated as coils [17, 21]. Following this assignment, the SS is segmented based on element types. Non-coil elements shorter than 2 residues in length are reclassified as coils, which are then excluded. The surviving helix and sheet elements undergo a transformation: helices are transferred into lower-case alphabetic characters, while sheets are converted into upper-case alphabetic characters (as depicted in Algorithm 1). For example E3 is translated into B and H2 is translated into a. Strings of these characters are termed Secondary Structure Elements (SSEs).

Proteins consisting of fewer than 5 SSEs are excluded. Moreover, as many residues folding into coils are discarded, our approach is tailored for proteins with a minimum of 50 residues for global comparisons, ensuring sufficient information for robust comparisons. Since protein domain size typically ranges from 50 to 200, the minimum number of residues was set as 50 for local alignment. [22]

#### Algorithm 1 Compressing Sheet and Helix Elements into Alphabetical Representations

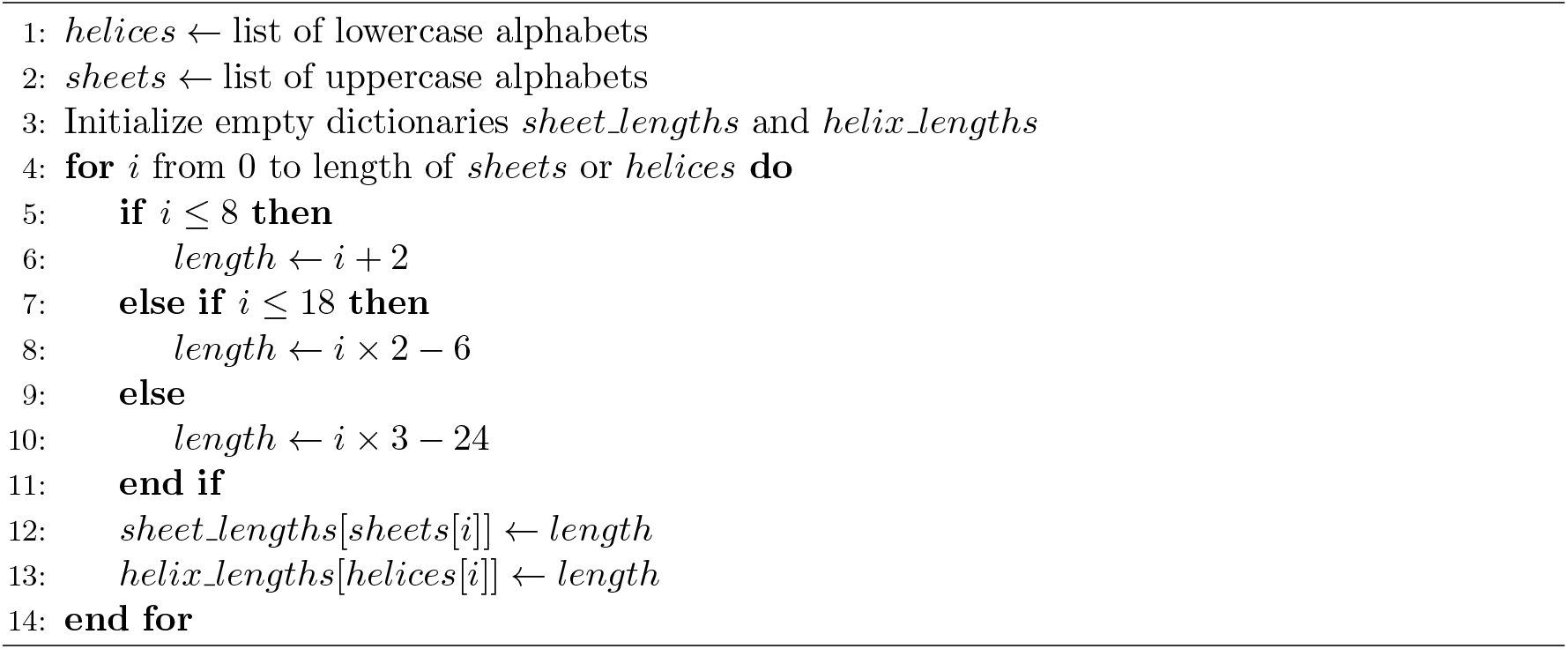

### 4.2 Similarity matrix for alignment

Instead of using the conventional binary paradigm for SS matching, we introduced a novel methodology involving the generation of customized similarity matrix specifically designed for the alignment of SSEs. These matrices are constructed through an optimisation process that uses a scoring mechanism to take both the type and the length of a given structural element into account. In addition, as the elements of SSE strings represent different numbers of actual residues, element-specific gapopen and gap-extension penalities were optimized for different elements, in contrast to the global gap penalties used in many of the standard sequence alignment algorithms.

The matrices enforce a scoring system in which aligned elements are scored higher for longer matches, while mismatches are penalized based on the difference in elemtent length. The smaller the length difference, the lower the penalty. Notably, diagonal elements in these matrices are assigned the highest scores within their respective rows and columns, ensuring that matches of the same length and type are optimally rewarded. Furthermore, the scoring can extend into negative values, with mismatches between different SSE types incurring penalties through matrix masking. Moreover, traditional sequence alignment applies consistent gap penalties against all the elements. By contrast, due to the compression, we employ a gap penalty that increases with the number of residues that the corresponding element compresses. This approach emphasizes the importance of both length and element type in determining alignment scores.

To optimize the parameters of these matrices, the Jaccard index between the extraction region and SCOPe domains were defined as the fitness, which was then optimized through a genetic algorithm [23], and the constraint was enforced on the whole population during this process. Since the SCOPe database mainly consists of single-domain proteins, we avoided the bias toward full alignment extraction by specifically selecting those pairs that have at most one shared domain between them and of which at least one protein is multi-domain. Using the substitution matrix optimized by this method, we achieved a Jaccard index of 0.794 for single intersection multi-domain proteins and overall 0.922 for all the SCOPe pairs.

### 4.3 Pre-filtration for alignment and ML predictions

Although the SSE encoding provides a compact, low-dimensional representation of protein structure, performing local alignments exhaustively across a large database scales as *O*(*N* ^2^) and yields mostly spurious hits. We therefore introduce a two-stage pre-filter that reduces both the alignment workload and the subsequent ML prediction cost, while preserving high sensitivity.

**1. K-mer matching with spaced seeds**

- Extract all length-*k* substrings (*k* = 3) from each SSE sequence, and generate “spaced seeds” using the pattern [1, 0, 1, 0, 1].
- Allow up to ±3 shifts in the structural-alphabet index at each seed position (i.e. any letter within ±3 code points is considered a match).
- **Local consistency:** Each diagonal requires at least max(3, ⌊*L/*3⌋) consistent seed-matches
- **Global threshold:** Candidates must accumulate ≥ max(5, ⌊*L/*2⌋) matched positions
- **Result:** Eliminates approximately 90% of candidate alignments while retaining 86.1% sensitivity for true TM-scores ≥ 0.6, and 90.1% sensitivity for predicted TM-scores ≥ 0.6.

**2. Normalized local-alignment score filter**

- For each remaining candidate pair (*A, B*), compute the local Smith–Waterman score SW(*A, B*) using our SSE-substitution matrix.
- Compute each sequence’s self-alignment score SW(*A, A*) and SW(*B, B*) based on the aligned region of (A,B), and define the normalized score

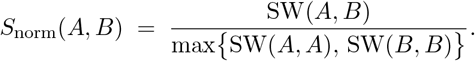

- Discard all pairs with *S*_norm_(*A, B*) below a chosen threshold.
- **Result:** Further reduces the set of ML predictions by 95%, while maintaining 89% sensitivity for predicted TM-scores ≥ 0.6. and 80% sensitivity for real TM-scores

### 4.4 Neural Network for local structure comparisons

Secondary Structure Elements (SSEs) alignments are transformed into a 3D tensor with dimensions (2,L,3), where:

- 2 represents the two sequences involved in the alignment,
- L refers to the total alignment length, doubled because the distance between each element in the alignment is also considered.
- 3 encodes three key features: the presence of helices, sheets, and the residues distances between the neighbouring SSEs.

This 3D representation encapsulates both the structural and distance-based relationships between aligned sequences, providing a rich data structure for learning. (A graphical representation is shown in Figure 1c.))

To analyse this data, we designed a hybrid neural network architecture that integrates a 2D ResNet [24] and a bidirectional Long Short-Term Memory (BiLSTM) model [25]. The ResNet component is responsible for extracting spatial features from the 3D tensor by processing the paired SSE sequences in a two-dimensional convolutional manner. Simultaneously, the BiLSTM model captures sequential dependencies and patterns within the alignment data by operating along the length dimension, L, effectively learning both forward and backward temporal patterns.

This hybrid model is designed to leverage the strengths of both CNNs (for alignment pattern recognition) and BiLSTM (for sequential pattern analysis), providing an effective approach for learning complex patterns in SSEs-based alignments.

### 4.5 Domain boundary extraction from alignment

In our method, we summarize domain boundaries from large-scale pairwise alignments using Density-Based Spatial Clustering of Applications with Noise (DBSCAN) [26]. DBSCAN identifies clusters based on density, where core points are defined as having at least *n* neighbours within a distance *ϵ*, as shown in Equation 1. If a point is not reachable from any core point, it is considered noise and excluded from the clustering process. This noise exclusion plays a crucial role in our dataset, where small variations in domain boundaries occur frequently, and removing these noisy points helps to improve the precision of the clustering.

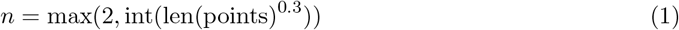

The distance threshold *ϵ* is determined using the Interquartile Range (IQR) method [27]. This method helps adjust the clustering process to the variability in domain boundary positions and provides a robust measure against outliers.

By applying this IQR-based threshold, the DBSCAN algorithm effectively handles datasets with diverse domain sizes and levels of redundancy. This method allows the clustering process to group similar boundaries together while excluding outliers that could otherwise skew the results. As a result, the algorithm achieves a better balance between sensitivity to local variations and robustness to noise, making it well-suited for large-scale alignment analysis.

### 4.6 Space Possibility and Information Content

The original sequence and 3Di sequence both consist of 20 elements, giving a total number of possible sequences:

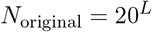

After compression, each position in the SSEs has 52 choices, but the length is reduced to approximately 7.7% 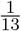 (see Figure 2). The space possibility for SSEs is:

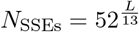

The compression ratio of the sequence space is then:

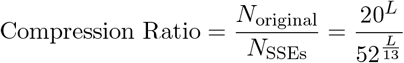

Simplifying using logarithms:

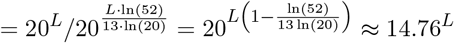

From an information-theoretic perspective, the information content of a sequence can be calculated using Shannon entropy [28]:

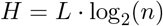

For the original sequence:

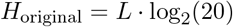

And for the compressed SSEs sequence:

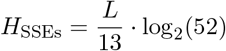

The compression efficiency, representing how much the information content is reduced, is:

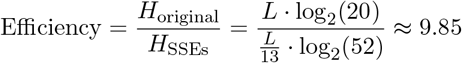

### 4.7 Foldseek

Two similarity metrics in FoldSeek (FS) were compared: (1) 3Di sequence identity and (2) probabilities of SCOPe superfamily membership. This comparison aimed to identify which metric better reflects structural similarity through F1-score analysis. As demonstrated in Figure 8, the probability-based metric substantially outperforms 3Di sequence identity.

**Figure 8.**
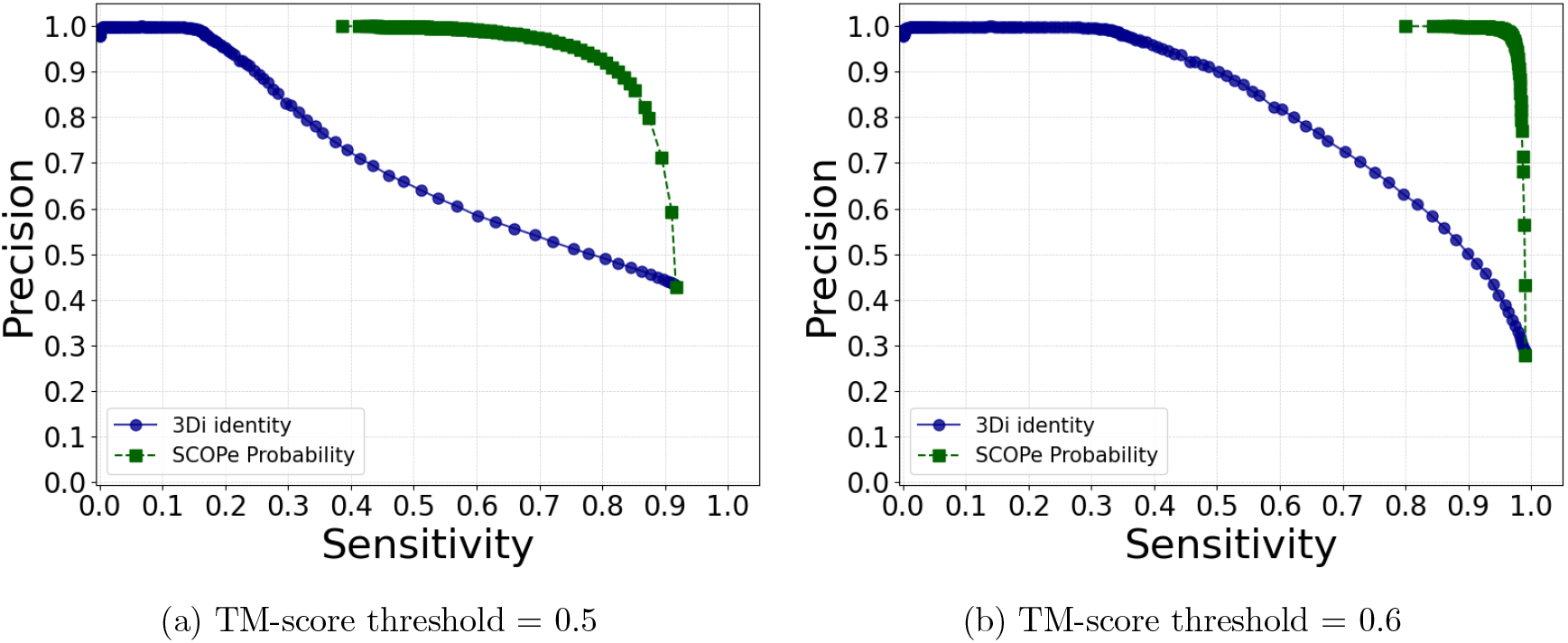
Sensitivity-Precision based on 3Di sequence identity and probability of being a SCOP domain. Clearly, using the probability index can better preserve high structurally similar pairs with high confidence. Using TM-score=0.6 as threshold will have a slightly higher sensitivity and higher precision. Therefore, TM-score threshold of 0.6 are used in later study.

FS’s local structural matches (Smith-Waterman [13]) were validated through dual TM-score analysis: global similarity (entire structure) and local similarity (aligned regions). Proteins with matching global/local TM-scores ≤0.6 confirmed true positives. For conflicting cases (FS match but global TM*<*0.6), local TM-scores of aligned substructures distinguished partial similarities from false positives. Length normalization (≈100 residues) controlled for size-related scoring biases.

A dataset of 5,000 non-redundant proteins was evaluated at TM-score thresholds of 0.5 andWhile TM-align defines TM-score ≥0.5 as structurally similar, FS exhibited reduced reliability at this threshold compared to TM-score ≥0.6 (Figure 8), establishing the latter as a more robust benchmark.

The probability metric demonstrated superior performance over 3Di sequence identity in identifying high-confidence structural matches, especially for distantly related proteins. Consequently, FS’s probability scores were adopted as the primary indicator for detecting similar substructures in subsequent analyses.

## Code Availability

The complete source code for the SSE-search pipeline, including scripts for SSE extraction, database construction, and structural alignment, is available on GitHub: https://github.com/rl647/SSE_search.

## Notes

https://github.com/rl647/SSE_search.

### Competing Interest Statement

The authors have declared no competing interest.

